# Characterization of Animal Movement Patterns using Information Theory: a Primer

**DOI:** 10.1101/311241

**Authors:** Kehinde Owoeye, Mirco Musolesi, Stephen Hailes

## Abstract

Understanding the movement patterns of animals across different spatio-temporal scales, conditions, habitats and contexts is becoming increasingly important for addressing a series of questions in animal behaviour studies, such as mapping migration routes, evaluating resource use, modelling epidemic spreading in a population, developing strategies for animal conservation as well as understanding several emerging patterns related to feeding, growth and reproduction. In recent times, information theory has been successfully applied in several fields of science, in particular for understanding the dynamics of complex systems and characterizing adaptive social systems, such as dynamics of entities as individuals and as part of groups.

In this paper, we describe a series of non-parametric information-theoretic measures that can be used to derive new insights about animal behaviour with a specific focus on movement patterns, namely Shannon entropy, Mutual information, Kullback-Leibler divergence and Kolmogorov complexity. In particular, we believe that the metrics presented in this paper can be used to formulate new hypotheses that can be verified potentially through a set of different observations and be complementary to existing techniques. We show how these measures can be used to characterize the movement patterns of several animals across different habitats and scales. Specifically, we show the effectiveness in using Shannon entropy to characterize the movement of sheep with Batten disease, mutual information to measure association in pigeons, Kullback-Leibler divergence to study the flights of Turkey vulture, and Kolmogorov complexity to find similarities in the movement patterns of animals across different scales and habitats. Finally, we discuss the limitations of these methods and we outline the challenges in this research area.

## Introduction

Information theory has always played an important role in biology [1, 2]. It is a field that is devoted to studying the storage, communication and quantification of information founded by Claude E. Shannon in his influential paper [3] and lies at the interface of mathematics, statistics, computer science and electrical engineering. While initial research in this field was mainly theoretical, we have witnessed a plethora of practical applications in the past decades. For example, concepts and techniques from this field have been used in several fields such as neurobiology [4], pattern recognition [5], cryptology [6], bioinformatics [7], quantum computing [8] and complex systems [9, 10] with significant success.

Recently, due to technological advances, low cost miniaturized sensors have been increasingly adopted for tracking the behaviour of animals across different scales and habitats. These sensors include, but are not limited to, Global Positioning System (GPS) receivers, accelerometers and radio-frequency identification (RFID) tags. This has led to an explosion in the deployment of these sensors in different habitats, across scales by animal behaviour researchers. A natural consequence of this development is that it is now possible to try to quantify and understand a variety of aspects related to animal behaviour such as migration patterns and routes, feeding, reproduction and mating patterns, conservation, monitoring of endangered species, epidemic spreading, resource use, social behaviour and association. In the field of animal behaviour, GPS, accelerometers and cameras are the predominant sensors deployed to measure several behavioural properties of animals. We limit our discussion here to GPS sensors due to their ease of deployment and the apparent simplicity of interpreting the data they produce as well as their consequent popularity relative to other sensors. For example, GPS sensors have been used to study selfish herd behaviour of sheep under threat [11], the hierarchical structures of group dynamics in flocks of pigeons [12], migration patterns in vultures [13], productivity in cows [14], and social relationships in birds [15] just to name a few.

We believe that information-theoretic approaches can provide *complementary* insights in the study of animal behaviour. In other words, these approaches do not replace the existing ones, but they are able to provide additional information about animal behaviour patterns, especially in terms of movement, which are not apparent using other types of analysis. Probabilistic approaches are also usually more robust in presence of noise, a common feature of sensor data. The number of applications of concepts and techniques from information theory to analysis of animal movement in the literature is limited. However, information-theoretic metrics have been used in the past for example to study information flow in animal-robot interactions [16] as well as predator-prey relationships [17, 18] in animals.

In this paper, we discuss how four classic information theoretic metrics, namely Shannon entropy [19], mutual information [20], Kullback-Leibler divergence [21] and Kolmogorov complexity (normalized compression distance) [22] can be effectively applied to the study of animal movement and provide additional *complementary* insights about animal behaviour. In other words, we explore how they can be used as tools for studying animal movement data. Indeed, the goal of this work is not to introduce new metrics, but to demonstrate the potential in using information theoretic concepts to understand animal behaviour. We introduce each metric separately and then we discuss how each metric can be applied to a practical problem, by discussing a case study in detail. More specifically, we demonstrate how these metrics can be used to characterize the movement patterns of animals across different scales and habitats. It is worth noting that these methods do not provide new ground-truth information, but they allow for identifying emergent patterns and formulating hypotheses that can be verified for example by means of further experimental observations in the field. The case studies are mostly based on datasets from the Movebank database [23].

The rest of the paper is organized as follows. In Section 1, we introduce the Shannon entropy as a method for the analysis of characteristics of movement and we demonstrate how it can be used to characterize the movement patterns of sheep with neurodegenerative disease. In Section 2, we describe the notion of mutual information and show how it can be used to measure association as well as reconstruct the flight dynamics in pigeons. We describe the Kullback-Leibler divergence in Section 3 and show how it can be used to characterize the annual movement patterns of the Turkey Vulture. In Section 4, we describe the Kolmogorov complexity (normalized compression distance) and demonstrate how it can be applied to extract the relationships in animal movement patterns across scales and habitats. We conclude our work by highlighting the challenges and a summary of the methods described in this work.

## 1 Shannon Entropy

### 1.1 Overview

The information content of a random variable is defined by the Shannon entropy [19] as a measure that quantifies the level of uncertainty embedded in such variable. In the case of animal movement, Shannon entropy provides a useful quantification of the level of regularity and predictability of the movement of an animal.

More formally, it can be defined as the uncertainty associated to a random variable *X* with realization *x*, which can be described by the following equation:

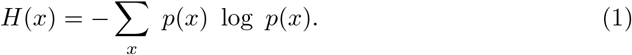

where *p*(*x*) is the probability density function and the summation is taken over all possible realizations of *X*. The base of the logarithm is not important and can take any value, provided the same base is used throughout the analysis.

Most of the information theoretic measures that exist today, some of which we will discuss below, are derived from Shannon entropy. However, there are other several definitions of entropy such as Rènyi entropy [24] and Tsallis entropy [25]. The discussion of these metrics is outside the scope of the present article.

Due to the noisy nature of most datasets, probabilistic metrics are becoming increasingly useful for modelling not only animal movement data but in general real world datasets to account for any form of uncertainty inherent in datasets of this nature such as missing data. Therefore, entropy can be used for assessing the overall welfare and well-being of animals instead of a metric like distance travelled. Most animals are known to have a regular activity-rest pattern except under highly unfavourable conditions or when they have some sort of impairment in their general well-being. This implies that animals are supposed to have a relatively high entropy except for the period of harsh conditions when they are either aestivating or hibernating. For this reason, entropy can be used to characterize the movement patterns of animals so as to assess the state of their health. In addition, it can also be used in lieu of tortuosity to describe how tortuous an animal’s path is using the turn angle as the input. The conditional entropy is an important element of the conditional mutual information and can be used for example in understanding swarm behaviour [26].

In the following subsection, we will consider a case study illustrating a possible application of Shannon entropy to the study of animal behaviour and, more specifically, to the characterization of the movement patterns of sheep with neurodegenerative diseases.

### 1.2 Case study: Shannon entropy as a tool for characterizing movement patterns of sheep with neurodegenerative disease

The detection of abnormal locomotion patterns is essential for the early diagnosis of a number of neurodegenerative diseases such as Batten disease in animals. Sheep with neurodegenerative diseases such as Batten disease are known to exhibit repetitive behaviours [27] over time due to gradual loss of motor skills [28] and social awareness [29]. Here, we use Shannon entropy to characterize the movement patterns of a flock of sheep comprising sheep with a natural mutation for Batten disease and their age matched control (mean age of 2 years) group using the dataset of [30]. We use the trajectory of each sheep sampled every second and compute the distance covered every ten minutes over eleven hours (20:00-7:00) each day for a total period of six days. This time window is chosen in order to minimize the influence of external environmental noise in the dataset. We further bin the resulting distance calculated in order to assign to each sub-intervals symbols. To bin the data, given that we are working with skewed distributions, we use the head/tail classification rule introduced in [31] resulting in 12 bins. First we evaluate this approach on synthetic data where we create two synthetic sheep sampling their distance covered from some random generalized Pareto distributions. We create ten of these distributions with different parameters. The abnormal sheep is allowed to sample from just two of these distributions while the normal sheep is allowed to sample from all. Details of this can be found in S1 Appendix. We compute the entropy and results Fig 1A) show the normal sheep have a higher entropy compared to the abnormal sheep. We further compute the entropy for each sheep (see Table 1 in S1 Appendix) as well as the mean entropy (Fig 1B) for the two groups of sheep. Our results show that the Batten sheep on the average have a lower entropy than the normal sheep with p-values (ANOVA) (0.0076, 0.1042, 0.2628, 0.0065, 0.0234, 0.0205) across the six days respectively. The potential impact of uncontrollable environmental variables such as unfavourable weather conditions is significant and may influence the behaviour of the sheep especially because the experiment was carried out in an open field. Therefore, the result should be interpreted with caution. We compare the entropy of the two groups of sheep with their respective average distance covered in (Fig 1C) and its mean variance (Fig 1D). The Batten sheep can be seen to have covered, on average, a longer distance over the period of observation.

**Table 1.**
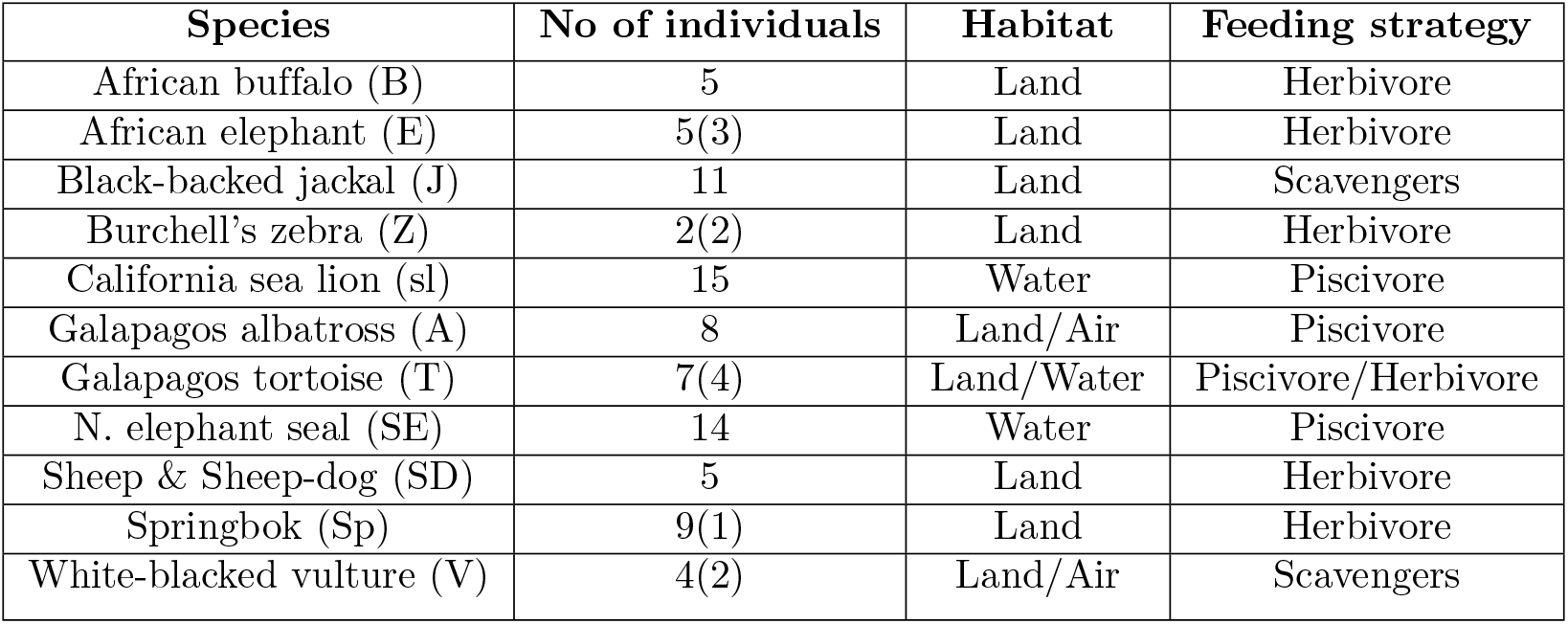
Summary of 85 individuals within 11 species used.

**Fig 1.**
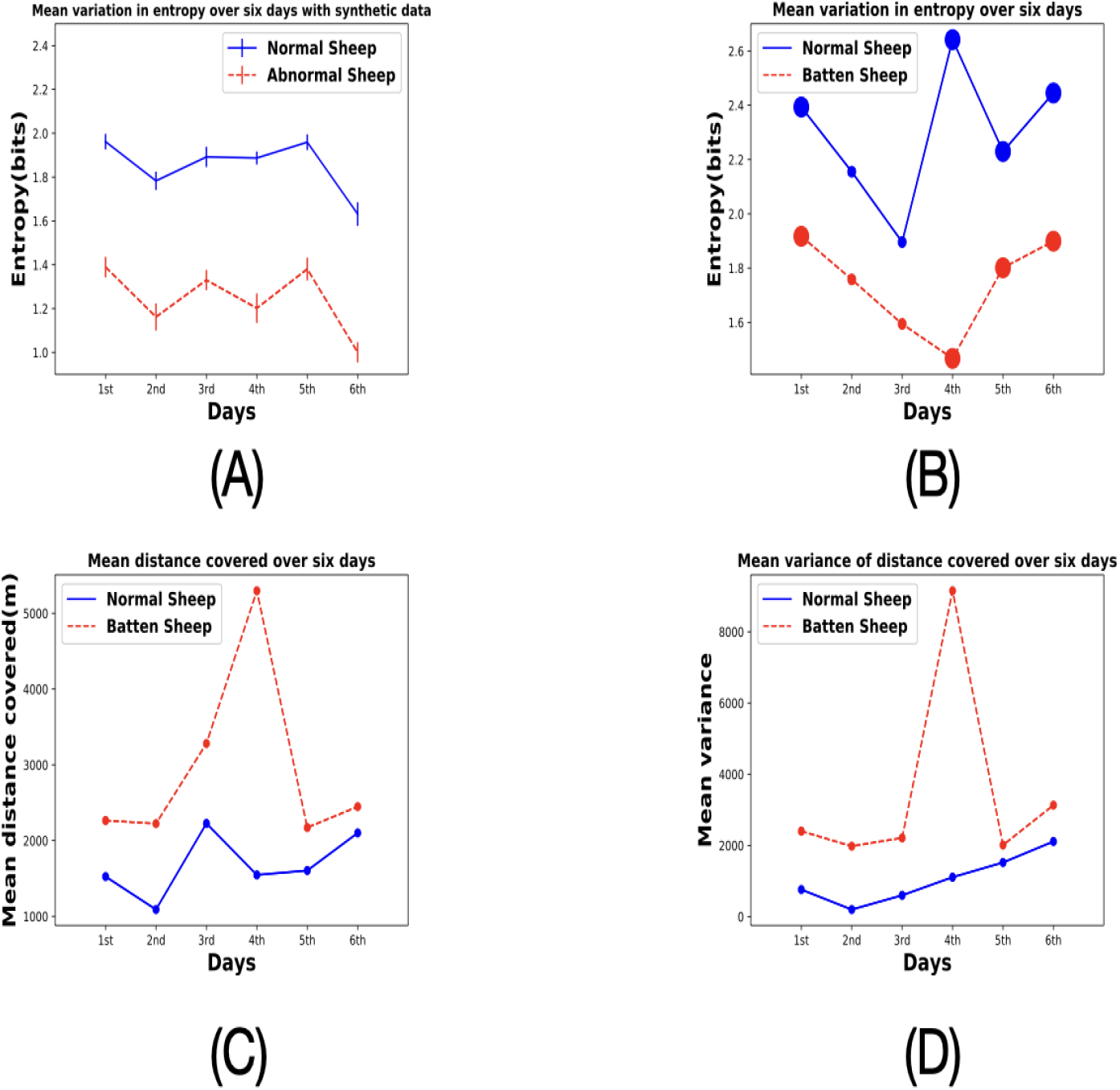
Entropy, distance and variance comparison. (A) Mean entropy with synthetic data; (B) Mean entropy of the two groups of sheep across 6 days where the larger circles represent days when the mean difference in entropy are statistically significant (One way ANOVA with p≤0.05). The sheep affected by Batten disease can be seen to have a lower entropy due to the tendency to repeat the same behaviour over a long period of time; (C) Mean distance covered by the two groups of sheep; (D) Corresponding variance of the mean distance covered.

## 2 Mutual Information

### 2.1 Overview

We now describe another classic information theoretic measure intimately linked to entropy, called mutual information [20]. The mutual information of two random variables *X* and *Y* defines the mutual influence one variable has over the other. Specifically, it quantifies the amount of information in one variable embedded in the other. For this reason, mutual information can be used as a measure of association or social-grouping, for example in the characterization of leader-follower relationship, group coordination and, more generally, collective behaviour [32]. This can be further used to derive a social network [33] and its complementary to the gambit of the group approach [34]. Mutual information can be used to measure non-linear relationships between two variables. More formally, the mutual information of two discrete random variables *X* and *Y*, with realizations *x* and *y* respectively, is given by:

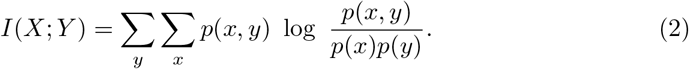

In this, *p*(*x*) and *p*(*y*) are the marginal probability distribution functions of *X* and *Y* respectively and *p*(*x, y*) is the joint probability distribution function of *X* and *Y*.

In the case of continuous random variables we have:

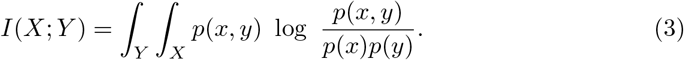

We now consider a potential application of mutual information, in its application to the study of association in pigeons.

### 2.2 Case study: Mutual information for measuring association and leadership in pigeons

As mentioned earlier, the de-facto method used in the animal behaviour community for measuring association is the gambit of the group otherwise known as co-location [34]. To detect significant associations and minimize co-location by chance, a permutation test (random shuffling of associations) is often carried out [35]. However, it is difficult to identify a general method for defining co-location. Also, the directional correlation delay time method (measures how long it takes for one bird to change direction relative to another) used by [12] in reconstructing pigeon flight network structure can only detect linear relationships leaving the non-linear relationships undetected. Previously in [36], the authors used transfer entropy (measured directed transfer of information) to infer leadership in Zebrafish. We state here that its our aim to detect association using a bidirectional graph and not a directed graph where information flow and its direction is of utmost importance. In addition, transfer entropy may be of limited use in instances where the agents are constantly changing positions relative to one another [37]. For a review of other methods that have been used for measuring leadership and influence, please refer to [38].

We demonstrate how Mutual Information can be used to overcome the limitations associated with the methods above by using it to measure association between pigeons in flight. We use the dataset of [37, 39] and select flight 8 as the result was discussed in the literature in detail. We compute the time-series of the turn angle of each bird followed by the pairwise mutual information of these time series of the nine birds involved in the flight to obtain a distance matrix (see Table 2 in S1 Appendix). As expected, there will always be a certain degree of association between all the birds in the flight, we use a randomization test to determine a threshold for significant pairwise mutual information (S1 Appendix). We further build a social network (Fig 2A) to visualize the flight formation. Our result is consistent with two previous studies on pigeon flight. First, we observe that pigeons do show a hierarchical formation when in flight as seen in (Fig 2A). This is a result consistent with the observations in [12]. Also, we were also able to detect the leader as node *M* during the flight which is the node *M* with one edge (Fig 2A)^1^ (see video of ground truth trajectory). This result is also consistent with the ground truth in the literature [37]. We compare this approach with three other methods: correlation coefficient, transfer entropy and Granger causality (Fig 2). While it is not straightforward to compare the performance of the four methods, one basis for comparison concerns leadership. Mutual information shows the best performance in terms of leader identification accuracy followed by the correlation coefficient. It is not possible to identify the leader using the transfer entropy and Granger causality. We attribute the poor performance of the transfer entropy and Granger causality to the continuous change in positions of the birds when flying.

**Fig 2.**
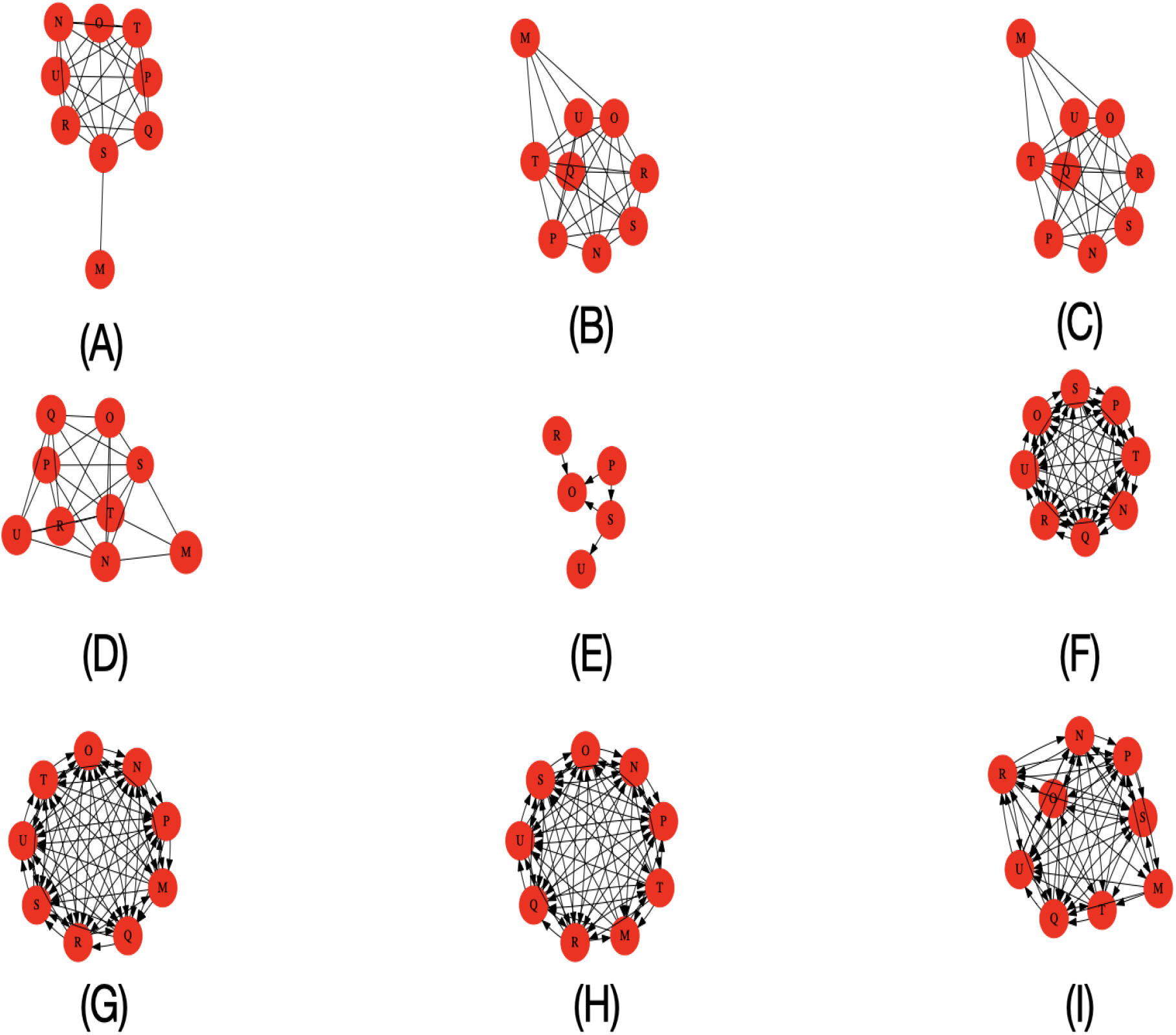
Flight dynamics of homing pigeons. Characterized by (A) Mutual Information(Kernel Estimator) [67] (B) Mutual Information(Discrete & Continuous mixtures) [68] (C) Mutual Information(Kraskov Estimator) [69, 70] (d) Correlation Coefficient (E) Transfer entropy (Kernel estimator) [70] (F) Transfer entropy (Kraskov estimator) [70] (G) Granger Causality (p-value = 0.05) (H) Granger Causality (p-value = 0.001) (I) Granger Causality (p-value = 0.0001). The tables presenting the pair-wise values can be found in S1 Appendix.

## 3 Kullback-Leibler Divergence

### 3.1 Overview

Kullback-Leibler divergence (KLD) also known as relative entropy measures the distance between two probability distributions. In animal movement modelling, KLD can be used to quantify changes in behaviour of an individual animal or discrepancies in behaviour in a group. For discrete probability distributions *P* and *Q*, KLD is defined as follows:

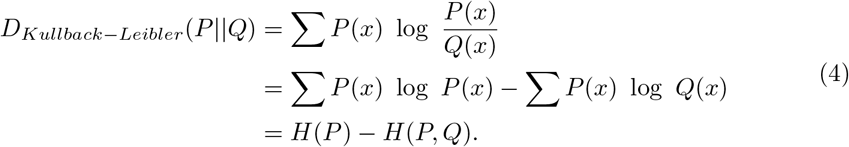

where *H*(*P, Q*) is the joint entropy between *P* and *Q* and *H*(*P*) the entropy of *P*. For continuous probability distributions *P* and *Q*, KLD is given by:

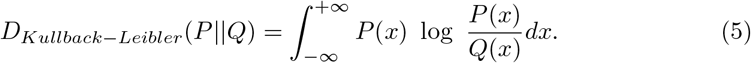

There are several potential applications of KLD. For example, it can be used for detecting behavioural change points and modes such as foraging, resting and travelling in animals by constructing a sliding window that moves across a time series while computing the KLD of the probability distributions of contiguous windows. This has implications for example in determining regime shifts, most especially for animals that move in non-homogeneous ways. In addition, it can also be used to identify points of change in landscape for animals who travel long distances over a heterogeneous landscape that affect their behavioural states. KLD has also applications in determining activity-rest patterns in animals.

Another potential application is the use of KLD for monitoring the health of animals that co-exist in groups. This can be achieved by computing the pairwise KLD between the probability distributions of movement data of all animals in the group while seeking animals with a significant divergence from the remaining members of the group. This approach, when integrated with appropriate machine learning tools such as hierarchical clustering, can be used potentially to classify animals into healthy and non-healthy or automatic classification of animals into species.

### 3.2 Case study: Kullback-Leibler divergence as measure por predictability of Turkey Vulture annual movement patterns

In this subsection, we demonstrate a potential use of KLD to a biological problem. KLD is indeed widely used as a measure for comparing probability distributions. Fore example, in [21] KLD is used to measure the divergence from the equilibrium behaviour of the blue tuna fish after a telemetry device was attached to it. In this case study, we use KLD to describe the movement patterns of the Turkey Vulture. The Turkey Vulture, is the world’s most abundant and widely distributed avian scavenger with a population in excess of five million individuals [40]. We characterize the movement patterns of the Turkey Vulture (*Cathartes aura*) [40, 41] across several years by comparing how the movement patterns at the beginning of the year (January) vary relative to the remaining months of the year across the next three years. The bird by name Leo was chosen since we have movement dataset for several years. We extract up to three years worth of movement data corresponding to 666 data points per month or 28 days and convert the coordinates to distance covered in kilometers. We compute the Kullback-Leibler divergence (symmetric version) across several years and our results (Fig 3) show that the movement strategy of this bird is highly predictable. We observe from Fig 3 that the Kullback-Leibler divergence across the three years is characterized by the presence of several peaks and troughs. The peaks represent the movement back to the breeding sites as well as the breeding season when there is little movement in the temperate regions of America. The troughs represent the migrating period when the birds migrate to tropical regions in search of food. These species of birds start breeding in the temperate regions such as North and South America where they have an abundance of food during the spring and this breeding continues until the onset of fall [40, 42]. In the colder seasons, these birds migrate to tropical regions where it is warmer and there is abundance of rain and food throughout the year. However, at the onset of spring around March, these birds migrate back to the temperate regions of America, where they are guaranteed abundant food and resources. We compare the KL divergence with the mean difference as well as the Earth Mover’s Distance (EMD) of the monthly movement data of interest (Fig 3). We also compare how the movement patterns in other months of the year (February to December) vary relative to the remainder of the dataset over three years. The results in Fig 4 show the bird have different movement patterns in the months between June and September relative to other months of the year, which is essentially its breeding season^2^.

**Fig 3.**
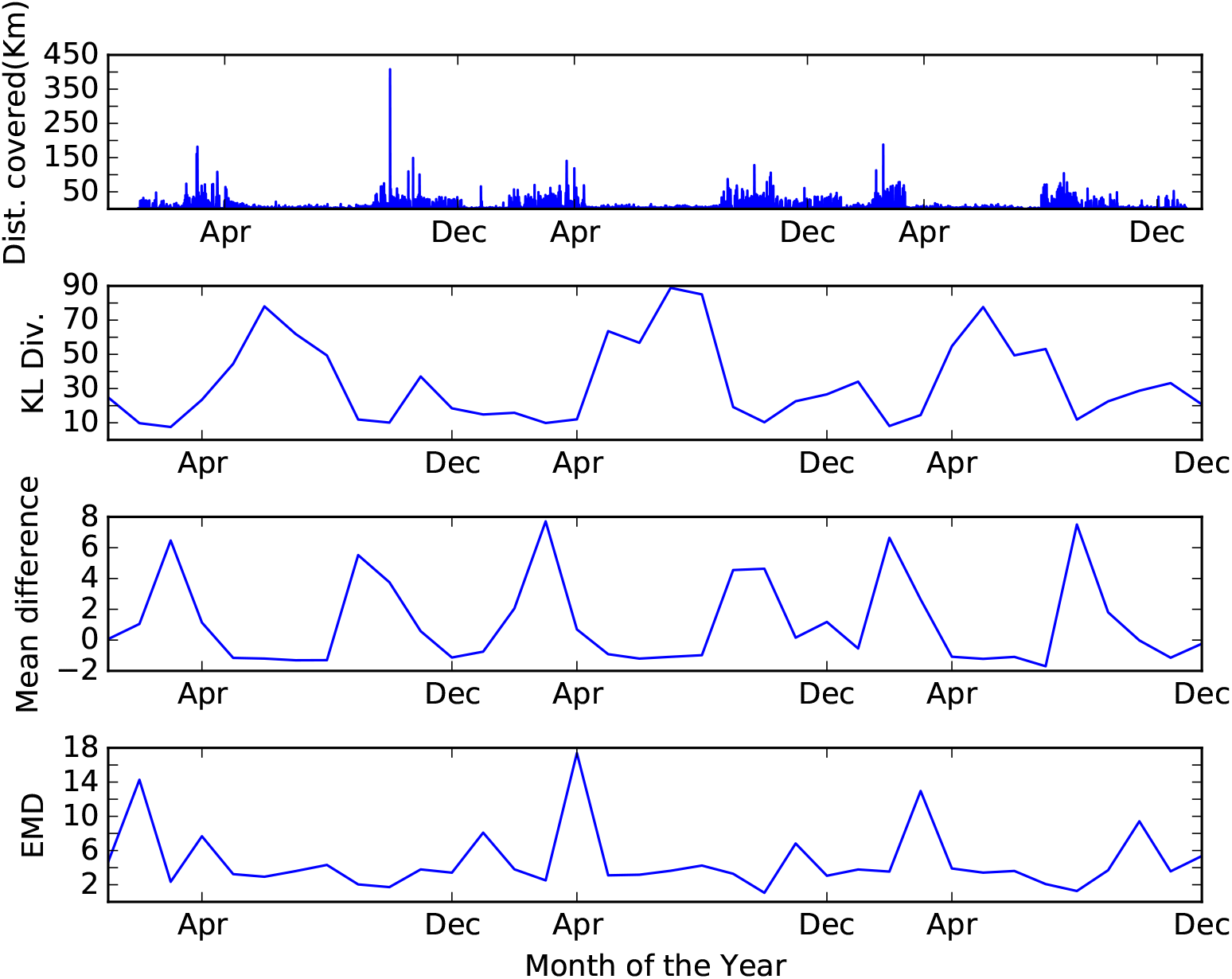
Kullback-Leibler divergence of the monthly movement pattern over a period of 3 years. The peaks represent the annual period of breeding as well as the flight back home after winter and the troughs migratory periods during which the bird travels in search of food. We compare the KL divergence with the mean difference and the Earth Movers Distance (EMD).

**Fig 4.**
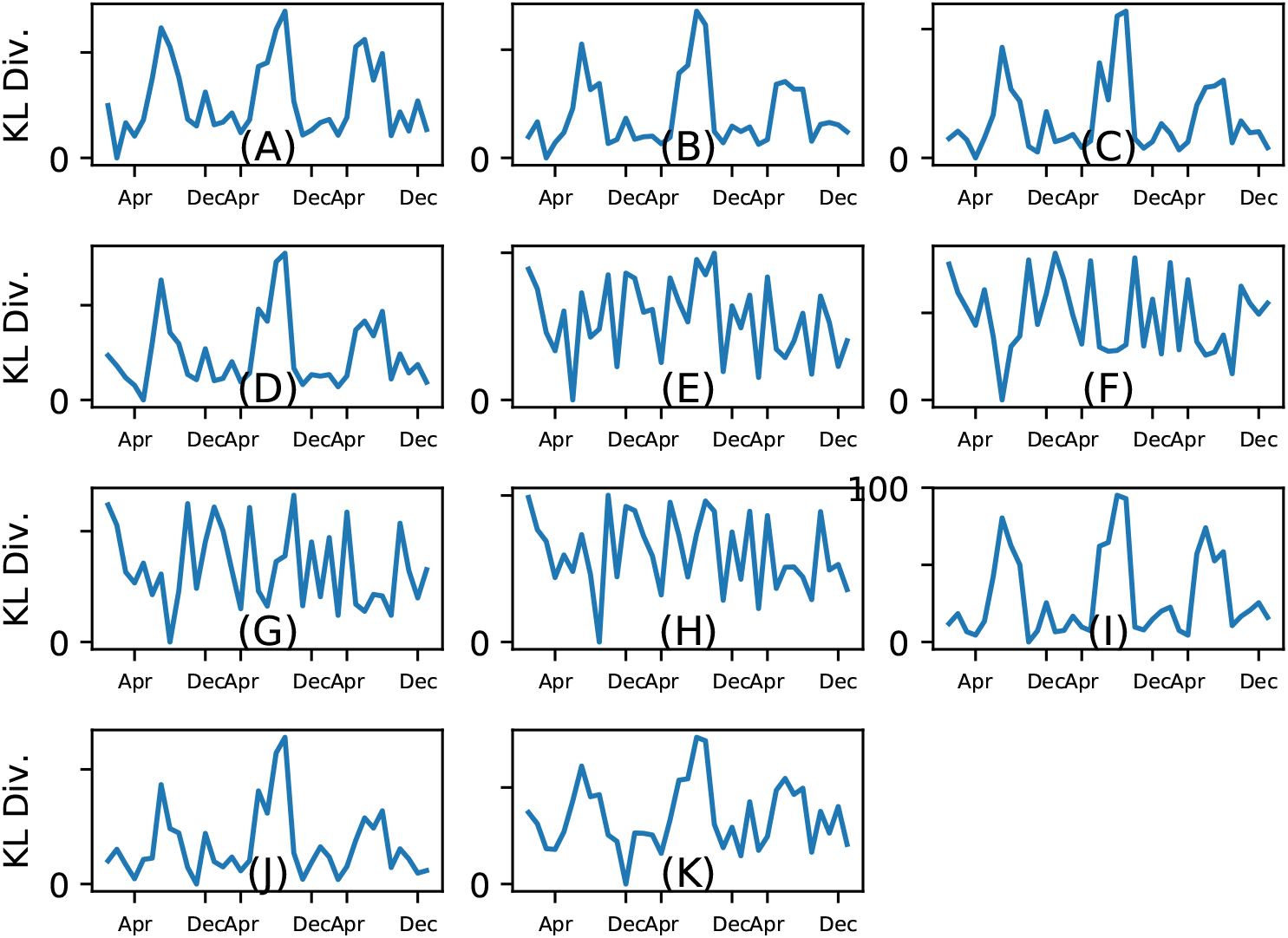
Kullback-Leibler divergence of the monthly movement patterns over a period of 3 years. Where the reference month is (A) February (B) March (C) April (D) May (E) June (F) July (G) August (H) September (I) October (J) November (K) December. From the result, it can be seen that the movement pattern during January, February, March, April, May, October, November and December are the same, while the bird exhibits a different pattern in June, July, August and September. This suggests that the breeding season of this bird is between June and September. The result is consistent with the information about Turkey Vulture in North America as they lay their eggs between May and June, incubate them for between 38 and 41 days and when the eggs hatch. The hatchlings are further brooded for a period between 70 and 80 days resulting in between 108 and 121 days of breeding, which is equivalent to four months of breeding [71].

## 4 Kolmogorov Complexity

### 4.1 Overview

Finally, we describe a similarity metric influenced by Kolmogorov complexity (KC), a metric with foundations in the field of algorithmic information theory. The Kolmogorov complexity of an object represents the shortest computer program that produces the object as output [22]. More formally, the KC of a string *x* with respect to a reference machine *U* is defined as:

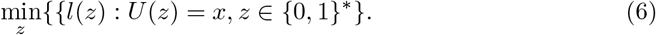

where *z* is a program that prints string *x* and then halts and *l* is the length. The concept of Kolmogorov complexity can be used as an inference tool to find the shortest description of behavioural data. The smaller the KC of a sequence the regular or simple it is and vice-versa. We describe the Normalized Information Distance (*NID*) [43], a similarity measure inspired by Kolmogorov complexity and defined as:

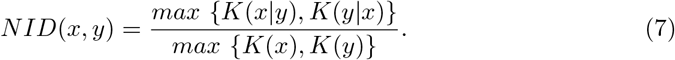

Due to the non-computability of the *NID*, the *NID* has been re-written [44] as the normalized compression distance (NCD) by simply approximating the Kolmogorov complexity *K*, using a compressor *Z*. The *NCD* between two strings *x* and *y* can be defined as:

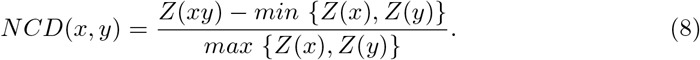

Here *xy* is the concatenation of x and y. These strings can be documents, software, genomes or even images. The *NCD* takes on non-negative values in the range 0 ≤ *r* ≤ 1 + *ϵ* with *ϵ* defined to take into account imperfections in the compression methods. Please refer to [44] for more details.

The *NCD* has been used in a variety of disciplines for different purposes, such as anomaly detection [45], gene expression dynamics [7], classification of music [46], detection and classification of computer worms and viruses as well as detecting the origin of new ones [47]. Since the *NCD* has been shown to work well with sequences and strings, it can be used, for example, in monitoring the behaviour of animals to know when they deviate from a previously or commonly known sequence of states, for example because of climate change [48]. It can also be used to quantify similarity in movement patterns of conspecifics across different habitats.

### 4.2 Case study: Kolmogorov complexity as tool for classifying animal movement patterns across scales

Animals across different habitats, scales, and species are known to have different movement patterns. However, little or no study has been carried out to find out which groups of animals possess similar movement strategies across different habitats and scales. Recently in [49], the authors discussed a classification of several animals across different species into similar groups using principal component analysis on some movement metrics with hierarchical clustering. Their result suggests that all animals organizes into four distinct groups of movement syndromes namely migratory, central place, nomadic and territorial. In this case study, we analyse the movement patterns of eleven animals (Table 1) across different spatio-temporal scales and habitats. For our analysis, we obtain the datasets of the Galapagos tortoise (*Geochelone nigra*) [50, 51], Springbok (*Antidorcas marsupialis*) [51, 52], African buffalo (*Syncerus caffer*) [51, 53–55], African elephant (*Loxodonta africana*)(original unpublished data contributed by Miriam Tsalyuk and Wayne M. Getz) [51], Black-backed jackal (*Canis mesomelas*) [51, 56], California sea lion (*Zalophus californianus*) (original unpublished data contributed by Dan Costa) [49, 51], Galapagos albatross (*Phoebastria irrorata*) [57], Sheep (*Ovies aries*) and Sheepdog [39, 58], Northern elephant seal (*Mirounga angustirostris*) [51, 59], White-backed vulture (*Gyps africanus*) [51, 60, 61] and Burchell’s Zebra (*Equus burchellii*) [62, 63]. All these datasets use the same 1 hour sampling period [49]. First, we evaluate our approach on synthetic data details of which can be found in S1 Appendix. Results, with both short and long movement data shows we can separate distinctly the synthetic animals into groups based on the distribution from which they sample their velocities. Then, we compare the monthly movement patterns of all the 85 animals to find similarities by computing their pairwise *NCD* with the gzip compressor [64] followed by hierarchical clustering of the resulting distance matrix (see S1 Appendix for details). The only metric used here is distance covered every hour, which we further processed to its binary equivalent (strings of zeroes and ones). We refrained from using the turn-angle here as it is an unreliable metric considering the noisy nature of most sensors. Our results (Fig 5) show three groups emerge: those that live on land (zebra, elephant, springbok, jackal, sheep and buffalo), those that live in water (tortoise, sea lion and elephant seal) and those that fly (Albatross and Turkey Vulture). Amongst the animals that live on land, we notice there appears to be some similarity between the movement patterns of the elephant and zebra while others seem to organize into distinct groups of conspecifics. Therefore, we hypothesize that there might be a correlation between the feeding and movement patterns of animals. We observe a small number of unexpected classifications: for example, Vulture *V*_1_ was classified among the animals that live in water. A possible explanation is related to the fact that the dataset is noisy. To find long-term similarities among animals movement patterns we compare approximately one year movement data of 16 animals (Table 1) across six different species. We selected these datasets due to their temporal length. Results (Fig 6) show that there might be some similarities in the long-term movement patterns of vultures and jackals which are both scavengers. This supports a hypothesis that there might be a correlation in the movement patterns of animals with similar feeding habits.

**Fig 5.**
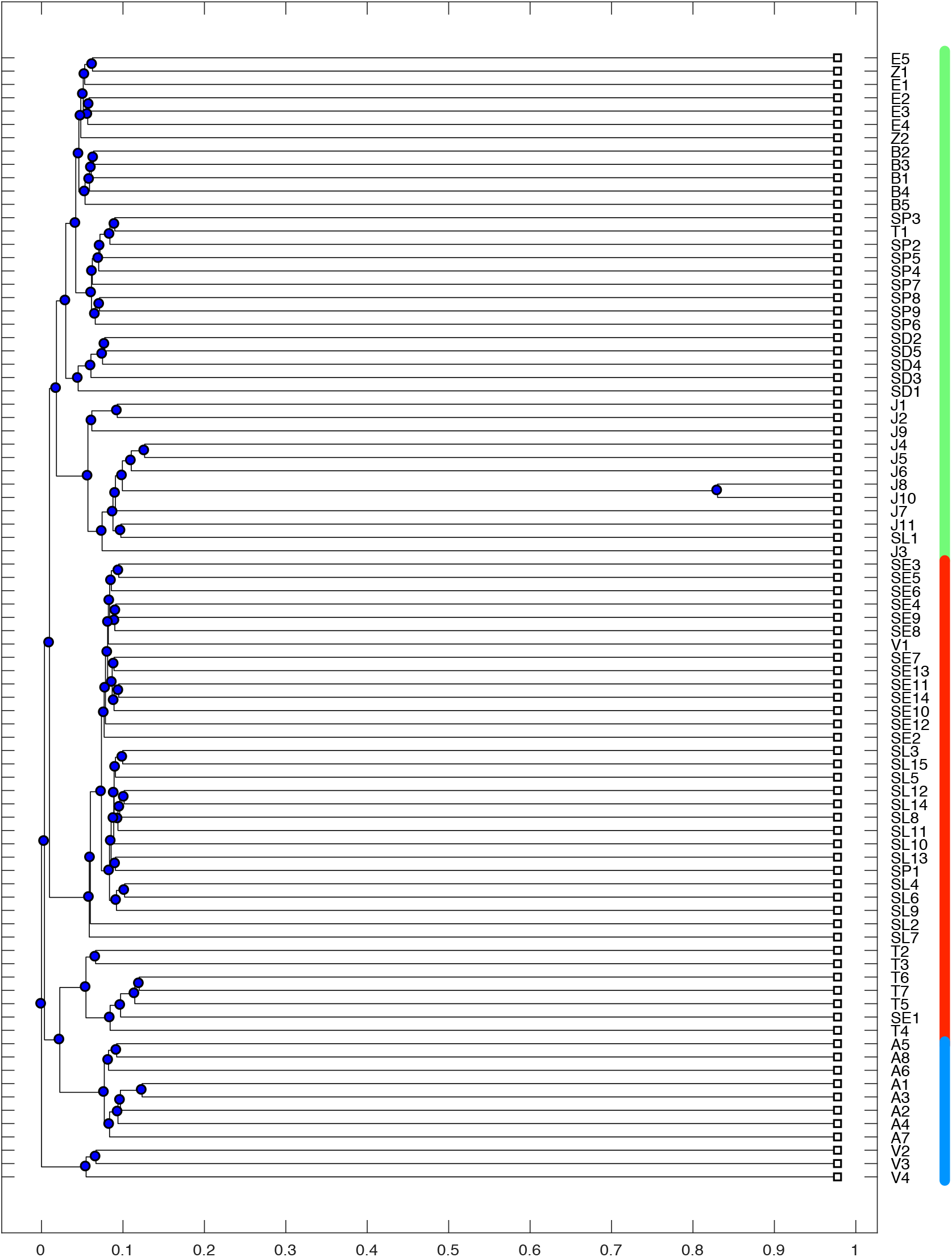
Hierarchical clustering of the pairwise NCD of 85 animals spread across 11 species. All the animals on average organize into three groups of those that live on land (green), those that live in water (red) as well as those who fly (blue).

**Fig 6.**
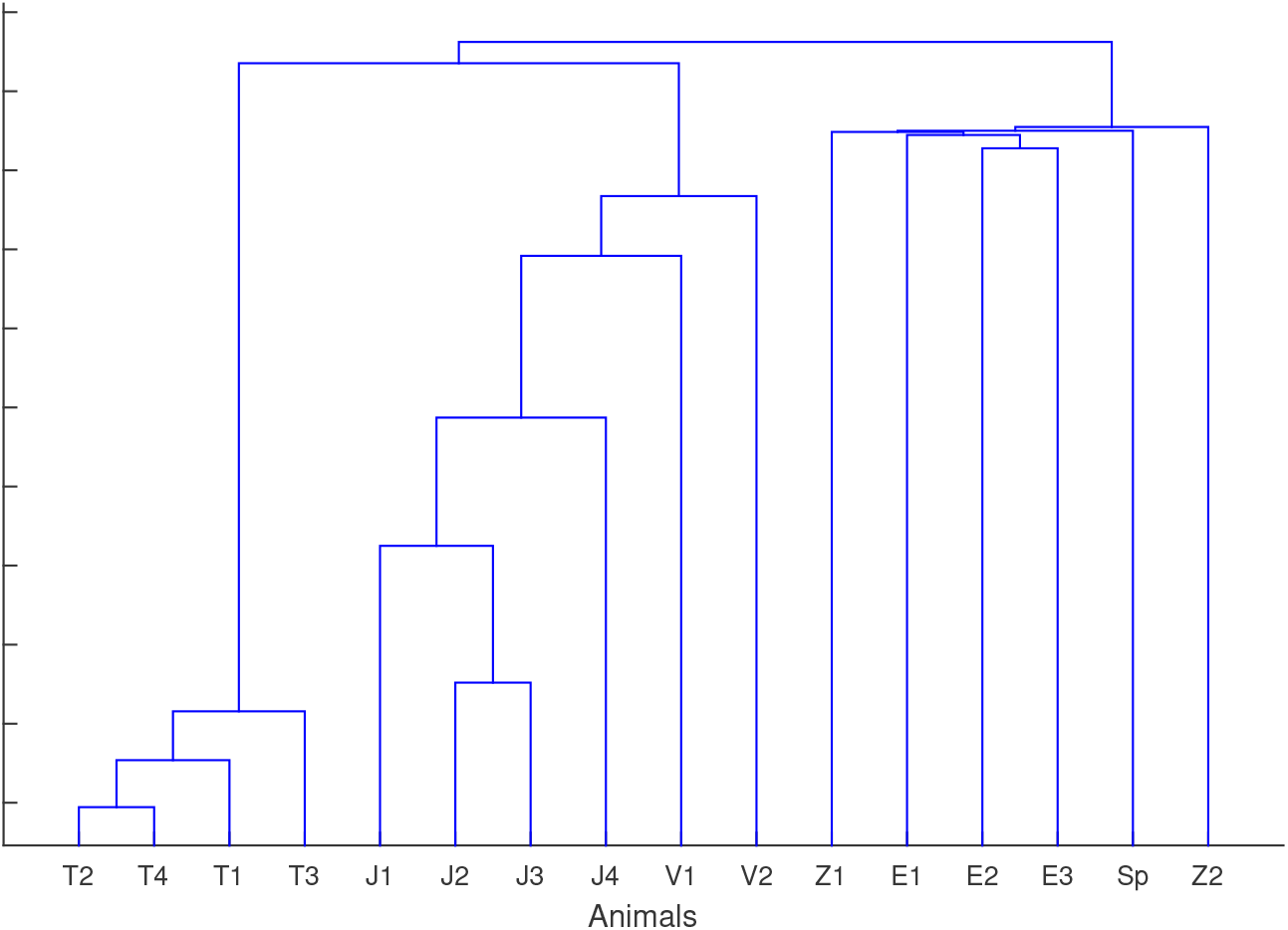
Hierarchical clustering of the pairwise NCD of 16 animals. These animals are spread across 6 species representing the movement patterns over a period of one year. The animals organize into three distinct groups that are correlated with their feeding patterns.

We compare this approach with the pairwise mutual information of the distance covered with respect to the two instances discussed above (see S1 Appendix). We believe that the insights obtained by means of this technique might be of value for researchers in order to formulate new research hypotheses based on the patterns emerging from this data.

The code in front of the species represents the label codes used for each animals in the hierarchical clustering while the number in the bracket under the no of individuals represents the number of animals with up to one year of observational data.

## 5 Limitations and Open Challenges

In this paper, we have highlighted the potential use of information theoretic metrics in obtaining insights about animal movement and we have also showed the applications of these metrics to real animal movement data. However, we would like to underline that these methods should be applied with caution given their inherent limitations. First, the issue of missing data remains a challenging problem due to logger failure or inability to regularly obtain position fixes. In addition, it suffices to state here that care must be taken while choosing the appropriate amount of data from which inference can be made. For example, let us consider the analysis of the similarities of the complexities of animals across different taxa and spatio-temporal scales. We identified similarity only in the long-term movement patterns of jackals and vultures. This phenomenon might not have been observed if datasets of shorter length had been used. Indeed the results derived from using the measures presented in this paper should always be evaluated considering the corresponding temporal scale of the dataset.

We issue a caveat on estimating probability density functions (PDF) of continuous movement data. At the moment, most of the methods and tools available are based on the assumption of underlying normal distributions. Considering that continuous animal movement data often follows skewed (e.g., power-law or truncated power-law) distributions [65, 66], researchers employing some of the methods described here, for example the Kullback-Leibler divergence, should exercise appropriate caution while estimating the PDF of these distributions. In the present study, we took special precaution while computing the probability distribution for entropy by binning the data around several mean values of the data in the direction of skewness.

Furthermore, appropriate methods for permutation and randomization must be used while determining a threshold, for example for the calculation of the pairwise mutual information between animals in a group.

## 6 Conclusions

In this work, we have demonstrated the use of a class of non-parametric information theoretic tools for studying movement patterns of animals and have showed how they can be applied by means of several animal movement datasets. First, we have demonstrated how Shannon entropy can be used to characterize the movement patterns of sheep with Batten disease where the distance covered every ten minutes was used as the basis for generating symbols to compute the entropy. The result shows that the Batten sheep have a lower entropy than their control counterparts. Then, we have described the use of mutual information for detecting associations in animals using pigeons as an example. Our findings show that this method can be very useful in lieu of the widely used gambit of the group approach as we were able to implicitly detect the leader from the flight data. We have showed how the Kullback-Leibler divergence can be used to characterize the movement patterns of the Turkey Vulture. From our results we were able to to evaluate the predictability of the Turkey vulture over time. Lastly, we have described a metric with foundations in the field of algorithmic information theory known as Kolmogorov complexity (normalized compression distance). We have used this metric to characterize the movement patterns of animals across different taxa and spatio-temporal scales with results suggesting there might be a correlation between the feeding and movement patterns of animals. These methods provide *complementary* insights in the study of animal behaviour. In particular, they can be used to formulate new hypotheses regarding animal movement. In other words, they provide additional information about animal movement that is not apparent using other types of analysis. This class of probabilistic methods is also usually more robust in presence of noise, which is inherent in location data.

As part of our future research agenda, we plan to explore additional movement datasets as they become publicly available and, possibly, other types of behavioural datasets, for example from accelerometers, in order to show further how these information theoretic metrics can be used to obtain novel insights about the behaviour of animals in their natural habitats.

## Supporting information

S1 Appendix (PDF)

## Acknowledgments

The authors would like to thank Professor Jenny Morton for her insight, valuable discussions, and for allowing them to make use of the sheep dataset in this paper.

## Author Contributions

**Conceptualization:** Kehinde Owoeye, Mirco Musolesi, Stephen Hailes

**Data Curation:** Kehinde Owoeye, Stephen Hailes

**Formal analysis:** Kehinde Owoeye

**Funding acquisition:** Kehinde Owoeye

**Investigation:** Kehinde Owoeye, Mirco Musolesi, Stephen Hailes

**Methodology:** Kehinde Owoeye, Mirco Musolesi, Stephen Hailes

**Project administration:** Mirco Musolesi, Stephen Hailes

**Resources:** Kehinde Owoeye, Stephen Hailes

**Software:** Kehinde Owoeye

**Supervision:** Mirco Musolesi, Stephen Hailes

**Validation:** Kehinde Owoeye

**Visualization:** Kehinde Owoeye

**Writing - original draft:** Kehinde Owoeye

**Writing - review & editing:** Kehinde Owoeye, Mirco Musolesi, Stephen Hailes

## Footnotes

1 https://www.youtube.com/watch?v=19srQkTVTGE

2 https://eol.org/pages/45511376/articles

